# Calculating the Mutual Information Between Two Spike Trains

**DOI:** 10.1101/423608

**Authors:** Conor Houghton

## Abstract

It is difficulty to estimate the mutual information between spike trains because established methods require more data than is usually available. Kozachenko-Leonenko estimators promise to solve this problem, but include a smoothing parameter which must be set. It is proposed here that the smoothing parameter can be selected by maximizing the estimated unbiased mutual information. This is tested on fictive data and shown to work very well.

## 1 Introduction

There are many problems in neuroscience that are addressed by examining the relationship between the spiking output of two neurons. The best way to do this should be to calculate the mutual information between the two spike trains: the mutual information is a measure of how much information the spike trains share and is therefore an ideal way of quantifying the relationship between them. However, in practice it has not been easy to use mutual information in this way because of the huge amount of data required to estimate it. This is because in the established method for calculating information-theory quantities for spike trains, the spike trains are converted into ‘words’ by discretizing time; this produces a huge number of words and so estimating their probabilities requires more electrophysiological data than it is typically practical to record.

In Houghton (2015) a Kozachenko-Leonenko estimator (Kozachenko and Leonenko, 1987; Victor, 2002; Kraskov et al., 2004; Tobin and Houghton, 2013) is presented for estimating the mutual information for random variables which take values on a metric space, that is, for data where there may be no coördinates, but where it is possible to measure the distance between two data points. This method is referred to here as the density estimation method. The density estimation method is relevant to the study of neuronal data because there are many metrics on spike trains (Victor and Purpura, 1996; Rossum, 2001; Aronov et al., 2003; Houghton and Sen, 2008; Houghton and Victor, 2010) which make the space of spike trains into a metric space.

This density estimation method for calculating the mutual information between spike trains relies on the choice of a smoothing parameter *h*. This paper describes an effective way to select this parameter and tests this method on fictive spike train data. It is found that the density estimation method produces a similar result to the more traditional binned method, but does so using considerably less spike train data.

The mutual information measures the dependence between two random variables *U* and *V* and is given by

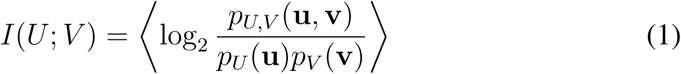

where 〈…〉 denotes the average with respect to the joint distribution *p*_*U*,*V*_ (**u**, **v**). **u** and **v** are values drawn from the *U* and *V* variables respectively. In the application considered here the aim is to estimate the mutual information between the activity of two neurons, so **u** and **v** are short intervals of spike train, one from each of the two neurons; in this paper 45 ms intervals are used. In order to calculate the mutual information the probability mass function *p*_*U*,*V*_ (**u**, **v**) needs to be estimated. The method for calculating mutual information described here is essentially a method of estimating the probability mass function.

The binned method for calculating the mutual information uses discretization. For a bin width *δt* each spike train interval is converted into a word by binning the spikes and counting the number of spikes in each bin. The mutual information is calculated on the words rather than the spike trains with the probability of a given word estimated by counting how often it occurs in the data. The advantage of this method is that in the limit of vanishing *δt* and of an infinite amount of data, the estimated mutual information approaches the true value. The disadvantage is that it approaches this true value very slowly. This is because of the huge number of words, for example, with 45 ms long spike train intervals and *δt* = 3 ms there are 2^15^ words and 2^30^, that is just over a billion, pairs of words corresponding to the spike train interval pairs. This situation can be improved with clever techniques (Treves and Panzeri, 1995; Nemenman et al., 2004; Magri et al., 2009) but one basic limitation is that it considers each word individually. The power of a Kozachenko-Leonenko approach is that is exploits the proximity structure of a metric space meaning the data points are considered in pairs.

Let 𝓟 = {(**u**_1_, **v**_1_), (**u**_2_, **v**_2_), …, (**u**_*n*_, **v**_*n*_)} denote a set of pairs of intervals from spike trains recorded during an experiment. These are modelled as being drawn from the probability distribution *p*_*U*,*V*_ (**u**, **v**). Following the formulation of the Kozachenko-Leonenko estimator given in Houghton (2015) this probability distribution is approximated by

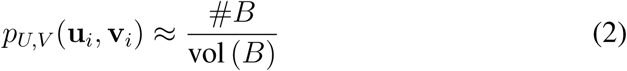

where *B* is a ball around the point (**u**_*i*_, **v**_*i*_), #*B* is the number of points in the ball and vol(*V*) is the volume of the ball. This approximation comes straight from the definition of the probability mass function

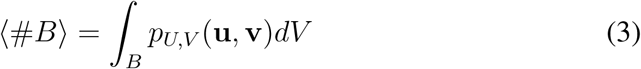

with the addition assumption that *p*_*U*,*V*_ (**u**, **v**) is approximately constant on *B*. The idea is that for every point a ball of fixed volume is used to estimate the probability mass function at that point. The volume chosen for this ball is a smoothing parameter: the larger the volume the more accurately #*B* estimates 〈#*B*〉, but for larger volumes the assumption that *p*_*U*,*V*_ (**u**, **v**) is approximately constant on *B* becomes less accurate.

The difficulty is how to calculate the volume of *B*. Because there are no useful coördinates for the space of spike trains there is no *dxdydz*-style integration measure. However, a probability mass function does provide a volume measure on a space and as described in detail in Houghton (2015) the marginalized distribution *p_U_* (**u**)*p_V_* (**v**) can be used to provide a volume measure on the space of spike train pairs.

Though this seems an odd choice of volume measure it gives a simple formula for the mutual information. For a point (*u_i_*, *v_i_*) consider the nearest *h U*-spike-train intervals to *u_i_*:

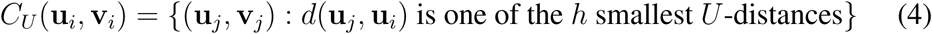

and the nearest *h V*-spike-train intervals to *v_i_*

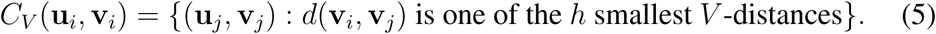

Now the ball around (**u**_*i*_, **v**_*i*_) is defined as

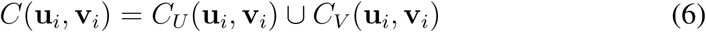

which has estimated volume *h*^2^/*n*^2^. Finally let #*C*(*u_i_*, *v_i_*) be the number of (*u_j_*, *v_j_*) points in *C*(**u**_*i*_, **v***_j_*):

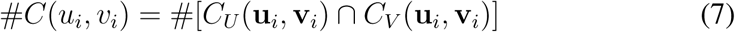

With this notation the Kozachenko-Leonenko approximation for the mutual information is given

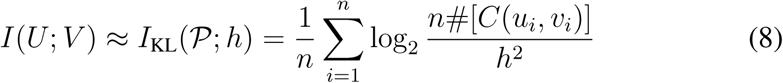

This quantity is straightforward to calculate. For each point (**u**_*i*_, **v**_*i*_), the set *C_U_*(**u**_*i*_, **v**_*i*_) contains (**u**_*i*_, **v**_*i*_) itself and the nearest *h* – 1 points to (**u**_*i*_, **v**_*i*_) when **u**_*i*_ is compared to the **u**_*j*_ in other (**u**_*j*_, **v***_j_*) pairs. Similarly, the set *C_V_*(**u**_*i*_, **v**_*i*_) contains the nearest *h* – 1 points when **v**_*i*_ is compared to *v_j_*. #*C*(**u**_*i*_, **v**_*i*_) is the size of the intersection. This is illustrated in Fig. 1.

**Figure 1:**
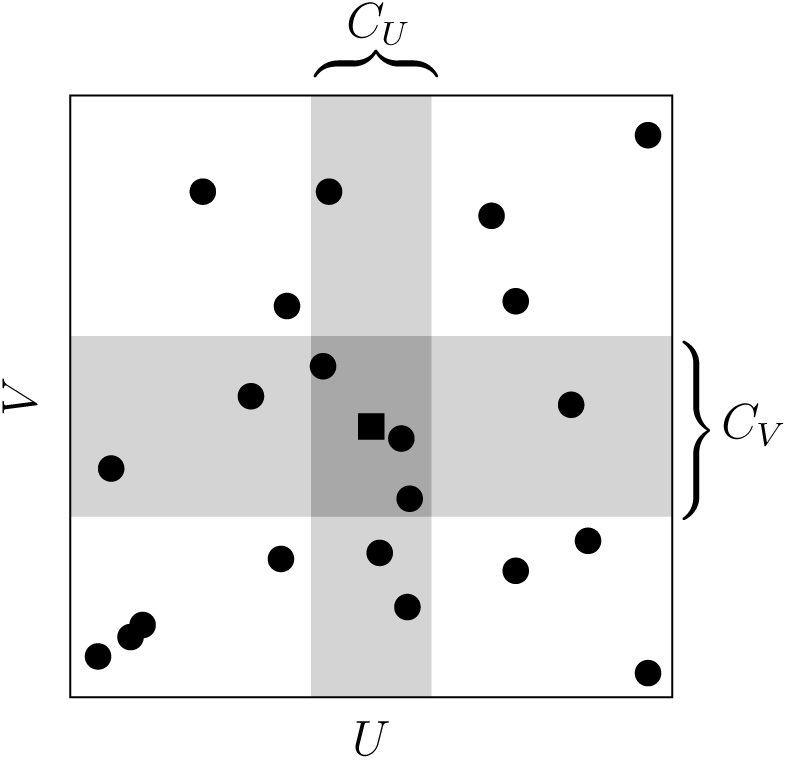
The calculation of *I*_KL_(𝓟; *h*). The *U*-space corresponds to the horizontal direction, the *V*-space to the vertical, of course, this is a cartoon, these spaces are not one-dimensional, they do not even have a defined dimension. The points in 𝓟 are marked as filled circles and a square; the square point is the one whose contribution to *I*_KL_(𝓟; *h*) is being calculated. The grey bars represent *C_U_*(■) and *C_V_*(■) with *h* = 7, that is, each of the grey rectangles contains seven points. #*C*(■) = 4 since it counts the points in the intersection, that is, the region with darker shading.

It is instructive to consider what happens if the two distributions are independent. In this case the it is possible to calculate the probability that #*C*(**u**_*i*_, **v**_*i*_) = *r* for different possible values *r*; it is a sort of urn problem. Choosing the *h* – 1 points in *C_U_*(**u**_*i*_, **v**_*i*_) that are not (**u**_*i*_, **v**_*i*_) itself is like randomly selecting *h* – 1 points out of *n* – 1 and calculating *r* is ask how many are in *C_V_*(**u**_*i*_, **v**_*i*_); this gives

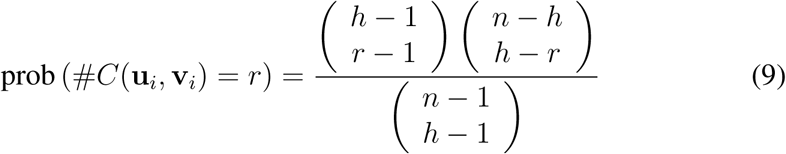

so

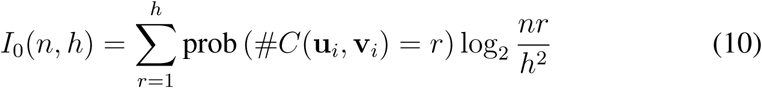

is the estimated mutual information when the two distributions are independent. This is a upward bias in the estimate of the mutual information; an upward bias is a common feature of estimators of mutual information. In this case the bias is because *B* will not always contain exactly 〈#*B*〉 points. One advantage of the Kozachenko-Leonenko approach is that *I*_0_ gives an explicit formula for the bias and it depends only on the smoothing parameter *h* and the number of pairs, *n*.

Obviously, as *h* approaches *n*, this bias approaches zero but otherwise it is positive and as bias it can be removed from the estimate of the mutual information:

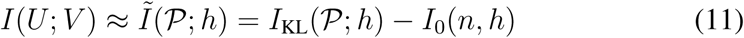

Recall that there are two competing approximations used in deriving the estimate: for small *h* the counting estimates for the number of points in a ball and for the volume of the balls are noisy; for large *h* the estimate of the probability mass function is too smooth. The first of these approximations is the cause of the bias described by *I*_0_(*n*, *h*). Conversely *I*_0_(*n*, *h*) is not effected by the smoothing bias. This suggests that the best approximation is found by maximizing *Ĩ*(𝓟; *h*) over *h*:

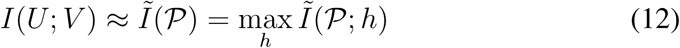

It is demonstrated here that this works very well.

## 2 Methods

### 2.1 Data

The algorithm is run on fictive data generated using two leaky integrate-and-fire neurons with shared input; the two neurons satisfy

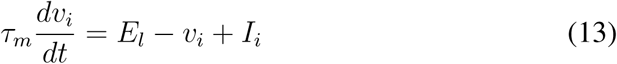

where, *i* = 1, 2 labels the two neurons, *τ_m_* = 12 ms and *E_l_* = –70 mV. If *v_i_* > *V_t_*, where *V_t_* = –70 mV a spike is recorded and *v_i_* is reset to *E_l_*; there is a refractory period of *τ_r_* = 2 ms. *I_i_* is an input, because the membrane resistance has been absorbed into *I_i_* it is a voltage rather than a current. Here

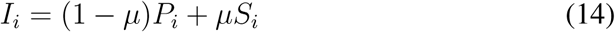

where *P_i_* is an input particular to the *i* neuron whereas *S_i_* is an input based on a shared input *S*:

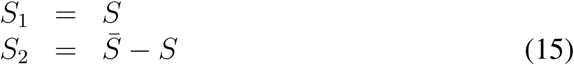

with *S̄* = 30 mV and the parameter *μ* specifying the amount of common input. The inputs *P_i_* and *S* are both piecewise constant, with each having a fixed value for a period chosen independently from a exponential distribution with mean *τ_c_* = 30 ms; the value for each interval is chosen uniformly from [0, *S̄*]. This network is illustrated in Fig. 2.

**Figure 2:**
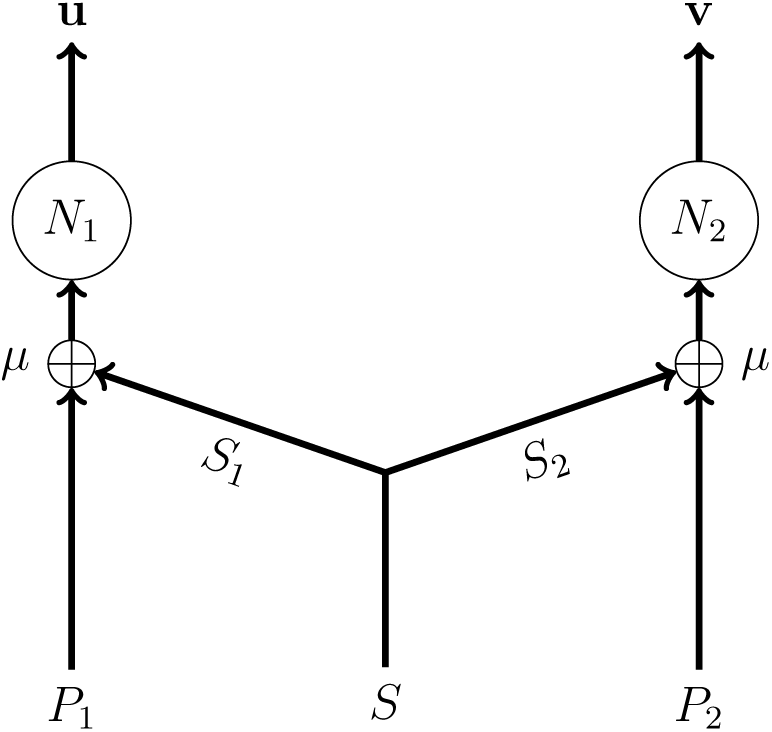
The fictive data. This shows the network used to produce the fictive data used to test the density estimation formula for mutual information. *N*_1_ and *N*_2_ are both leaky integrate-and-fire neurons, producing the spike trains **u** and **v** respectively. Their input is a weighted average of *P_i_* and *S_i_*; *S* is a shared input with *S*_1_ = *S* and *S*_2_ = *S̄* – *S*.

This method of producing fictive data is not perfect in the sense that the temporal correlation is different for different values of *μ* and the firing rate varies from 32 Hz at *μ* = 0 and *μ* = 1, to 27 Hz at *μ* = 0.5. However, the aim is to test the spike train pairs with different values of mutual information and, as will be seen, this method does succeed in doing that.

### 2.2 The distance between spike trains

For the density estimation information calculation the distance between individual spike trains is calculated using the van Rossum metric (Rossum, 2001), this calculates the distance between two spike trains **u** = (*u*_1_, *u*_2_, …, *u_n_*) and **v** = (*v*_1_, *v*_2_, …, *v_m_*) as

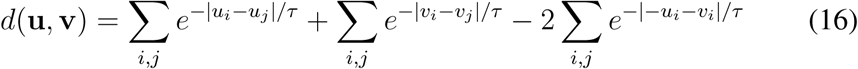

where *τ* is a time scale which can be thought of as expressing the precision of spike times in neuronal coding. A value around *τ* = 15 ms is often used. In the formula for mutual information the metric is used to order points by proximity so it might be expected that the values of estimated mutual information would not depend in a detailed way on the distance values or on the choice of metric. In fact, it will be seen here that in these data this holds true: the results are not sensitive to the value of *τ*.

### 2.3 Calculating the information

The pairs of spike trains produced by the network model are chopped up into 45 ms intervals. To calculate the mutual information using the binned method these intervals are discritised with *δt* = 3 ms giving 15 letter words. The frequency for each word pair is estimated from the data using the obvious empirical estimate

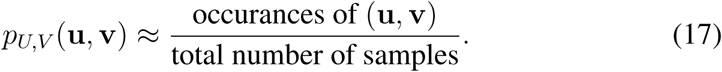

This is then used to calculate the mutual information directly. The bias is removed by also calculating the mutual information for shuffled data, in this case this means shuffling the pairing between the spike train intervals (Nirenberg et al., 2001; Montemurro et al., 2007; Panzeri et al., 2007; Magri et al., 2009).

To calculate the mutual information using the density estimation method, the distance matrix for the *U*-spike trains and the *V*-spike trains were calculated using the efficient implementation of the van Rossum metric described in (Houghton and Kreuz, 2012). The optimal value of *h* was found using a golden mean search (Kiefer, 1953).

## 3 Results

In Fig. 3 the value of the mutual information for 45 ms intervals of spike train is calculated for different values of *μ* ∈ [0, 1] using both the binned and density estimation approaches. For the density estimation approach 200 s of data is used; for the binned method 25000 s of data is used to establish the ground truth and smaller value of 2000 s to illustrate the amount of data needed to estimate the mutual information.

**Figure 3:**
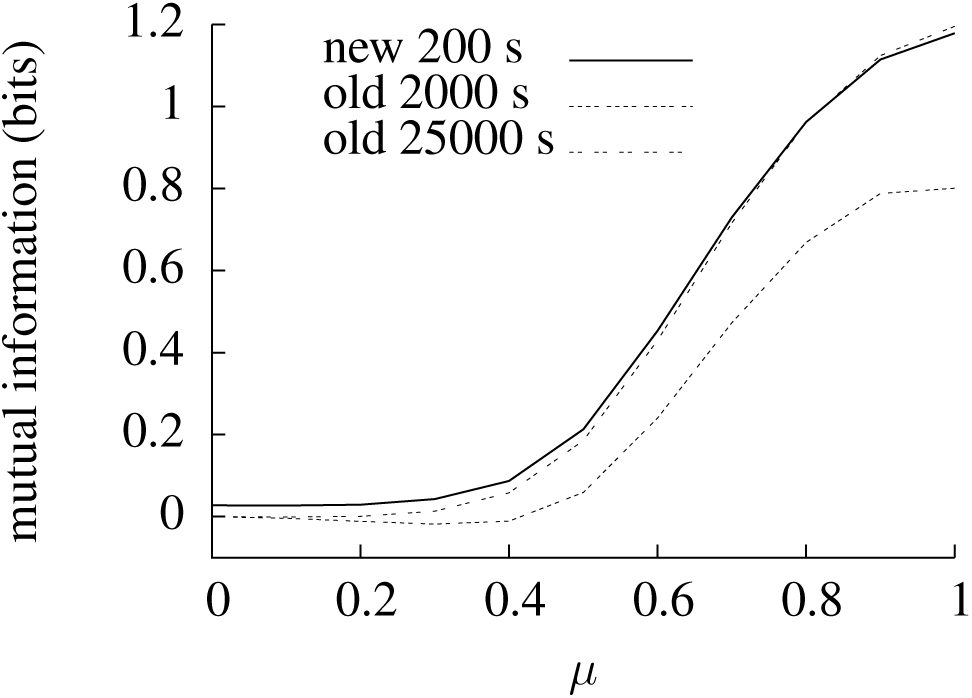
Estimates of the mutual information for different values of *μ*. The mutual information between pairs of 45 ms fragments of spike train is estimated using the density estimation, marked ‘new’ and the binned, marked ‘old’, approaches. In the case of the density estimation approach 200 s of spike train is used, in the binned approach, 2000 s and 25000 s are used. In all cases the graph shows the average of 100 trials; in the density estimation approach *τ* = 15 ms, in the binned approach 3 ms bins are used.

Using 2000 s of data the binned method gives a very poor estimate of the mutual information for most values of *μ*; the density estimation method is much closer to the value estimated using 25000 s of data. The binned method is better for values of *μ* < 0.4 when the amount of mutual information is very low; presumably this is because the noise in the estimate is more significant and the maximization over *h* leads to an over-estimate.

Fig. 4**A** and **B** show the convergence of the density estimation and binned methods; Fig. 4**C** uses a log scale to exhibit both on the same graph. These graphs show that the density estimation method uses considerably less data than the binned method. It is clear from these graphs that the estimators approach their asymptotic values in an orderly way. This means one approach, described in Treves and Panzeri (1995); Strong et al. (1998); Panzeri et al. (2007), to improving the estimate using the binned method is to fit the graph to a curve such as

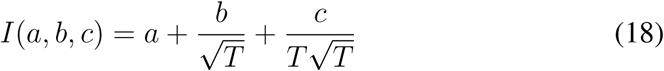

where *T* is the length of spike train used. For the simulated data being examined here this works quite well, concentrating on *μ* = 0.7 as an example, fixing *a*, *b* and *c* using the first 2000 s, gives an estimate of 0.6613 for *T* = 25000 s, compared to an actual value of 0.7156; the value given by the density estimation method using 400 s of data is 0.7412. The binned method gives even larger values if even larger amounts of data are used, for this value of *μ* 1000000 s of data gives 0.768413. Extrapolating the binned method from 200 s of data does not work, it gives an estimate of 0.1975. Fig. 4**D** compares the standard deviation for the two methods; they are roughly the same.

**Figure 4:**
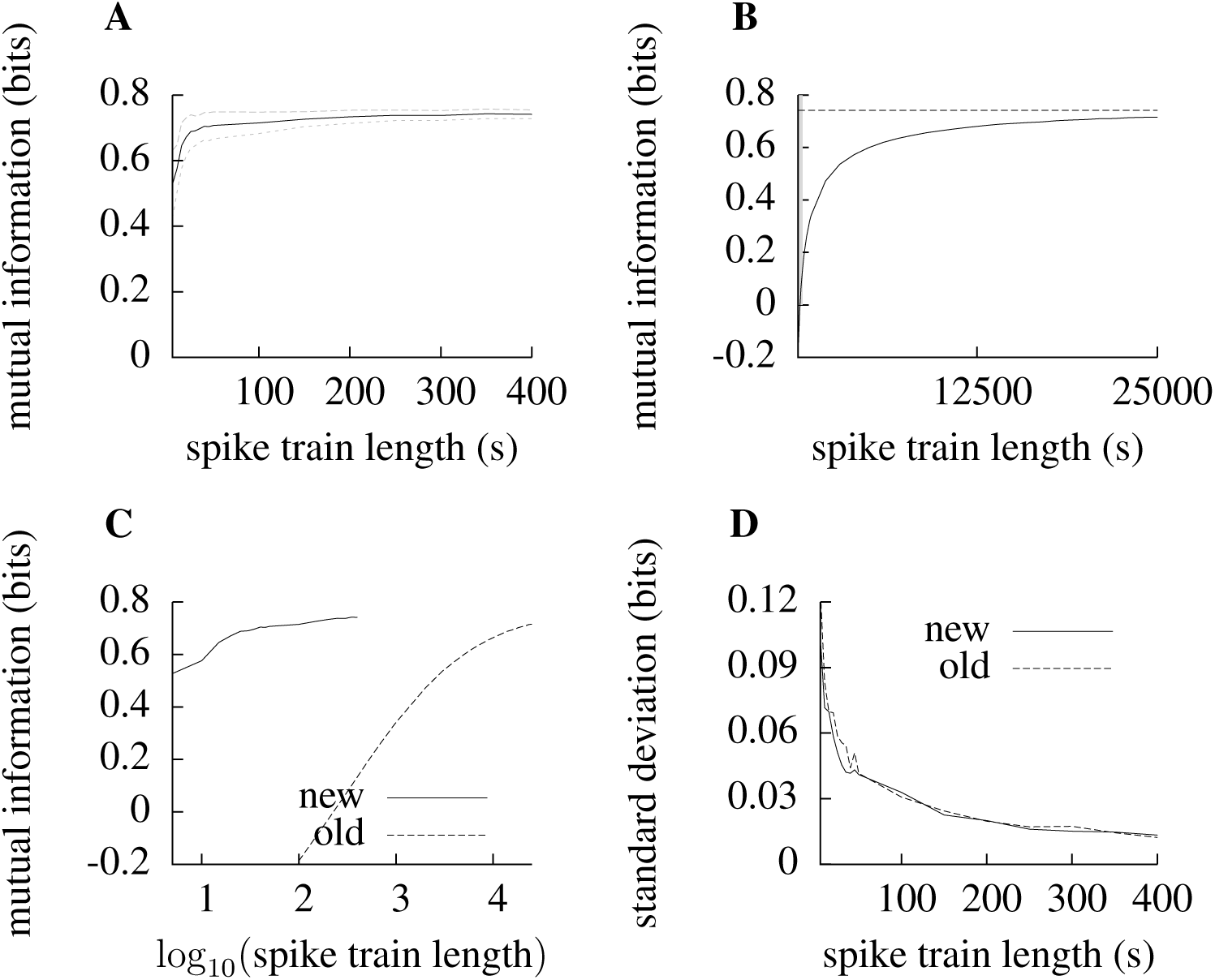
Performance of the formula for calculating mutual information. All these graphs concern pairs of spike trains where *μ* = 0.7 and the mutual information is being calculated between 45 ms fragments. **A** and **B** compare the density estimation (**A**) and binned (**B**) approaches to calculating mutual information as longer and longer spike trains are used. In **B** the very thin grey rectangle along the vertical axis marks out the area shown in **A** and the horizontal line gives the value estimated by the density estimation approach using 400 s spike trains. These two graphs are shown again in **C** where a log-scale is used for spike train length. In both **A** and **B** the plots are of the mean over 100 trials, in **A** the dotted lines show one standard deviation from the mean. For **B** the standard deviation is vanishingly small, this is because so much data is used in this approach. In **D** the standard deviation using the density estimation and binned approaches are compared, they are roughly similar, though, of course, in the binned approach the mean is very different from the value estimated using more data.

The robustness of the density estimation approach to mutual information is examined in Fig. 5. In Fig. 5**A** and Fig. 5**B** the lengths of the short intervals used to calculate the mutual information are changed; Fig. 5**A** also plots the binned estimate with a different letter length. In Fig. 5**C** a different stimulus is used, whereas for all the other simulations the shared input is shared with *S*_1_ = *S* and *S*_2_ = *S̄* – *S*, in this figure *S_i_* = *S* for both values of *i*. Finally, in Fig. 5**D** an input with a higher firing rate is used. The density estimator performs well, but there is some indication that the amount of data required, though modest compared to the binned approach, does increase as the number of spikes increases.

**Figure 5:**
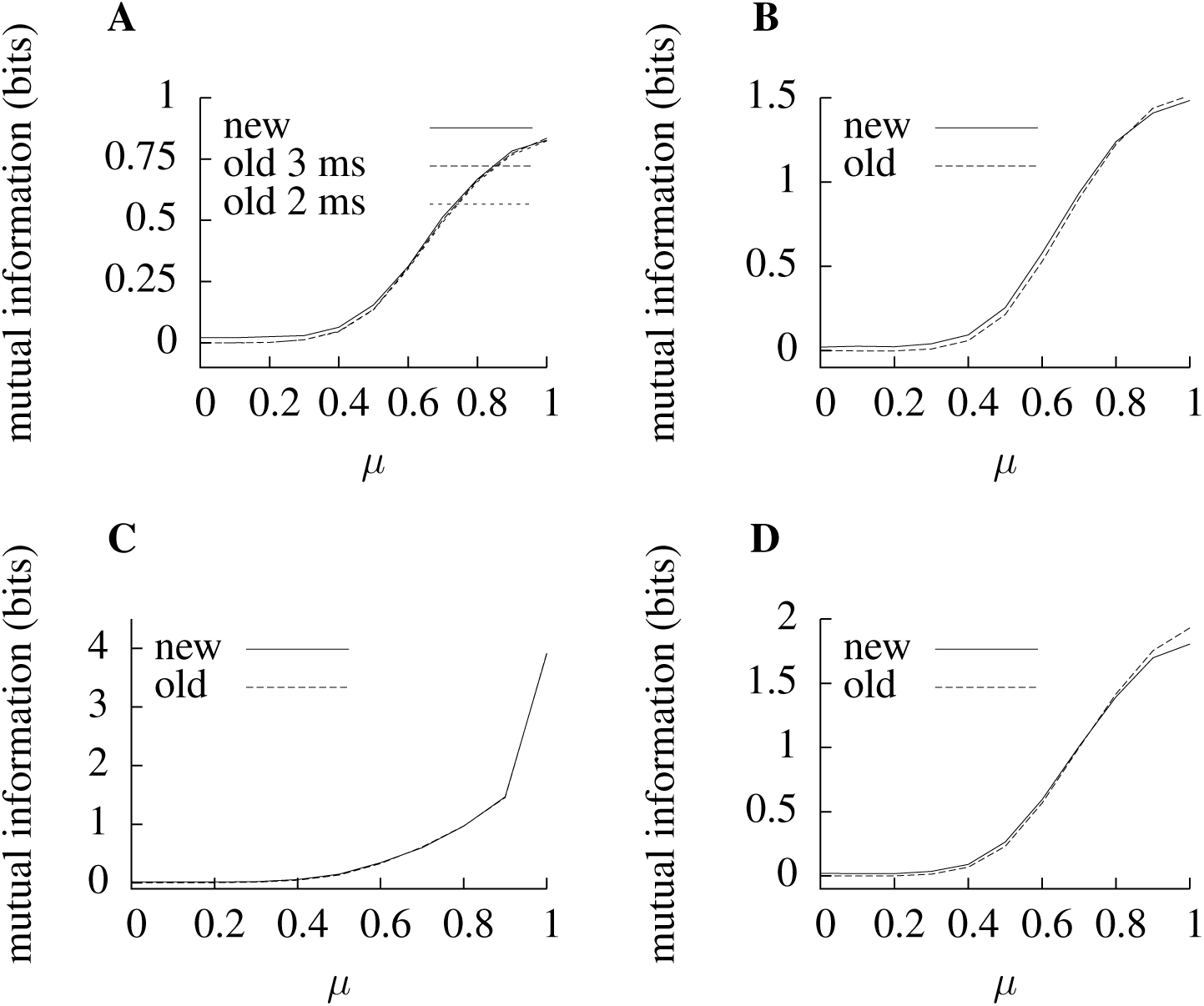
Testing the robustness of the density estimation approach. In **A** and **B** the length of the spike train intervals is changed. In **A** they are shortened to 30 ms, in **B** they are increased to 60 ms. In each case for the binned approach, marked ‘old’, the intervals are discretized into 3 ms letters, in **A**, the binned approach with 2 ms letters is also plotted. In **C** an alternative model is used to produce the simulated data: the shared input is the same, so the input is *S_i_* = *S* for both values of *i*. In **D** *S̄* = 35 mV is used to generate the simulated data, this increases the firing rate so that it varies from 44 Hz at *μ* = 0 and *μ* = 1 to 39 Hz at *μ* = 0.5. For the binned approach, 25000 s of spike trains are used in used in each case. For the density estimation approach, marked ‘new’, 200 s of spike trains are used in **A** and **C** and 500 s in **B** and **D**; when the intervals contain on average more spikes the density estimation approach appears to require more data. In **D** the difference between the two approaches is more noticeable for *μ* near one than in other graphs.

The sensitivity of the estimate to the choice of metric is examined in Fig. 6. In Fig. 6**A** the van Rossum metric is replaced by the Victor-Purpura metric (Victor and Purpura, 1996). This was the first metric proposed for spike trains and rivals the van Rossum metric in measures of how well metric distances capture information coding in spike trains (Houghton and Victor, 2010). Furthermore, it is a non-Euclidean metric (Aronov and Victor, 2004); the van Rossum metric works by embedding the space of spike trains into the infinite-dimensional space of functions, in this sense the van Rossum metric is Euclidean, replacing it with the Victor Purpura metric demonstrates that this Euclidean property is not required for the density estimation approach to work. In Fig. 6**B** small values of *τ* show a poor performance; for small values of *τ* the metric distance between two spike trains is very dependent on noise which jitters spike times to a degree that is significant when compared to *τ*; for values of *τ* that are similar or larger than the size of the interval, the relative distances between different pairs of spike train does not change as *τ* varies. The behaviour of the smoothing parameter is explored in Fig. 7; Fig. 7**A** shows an example of how the estimated mutual information depends on *h* and Fig. 7**B** graphs the change in the optimal value of *h* as the length of the spike trains changes.

**Figure 6:**
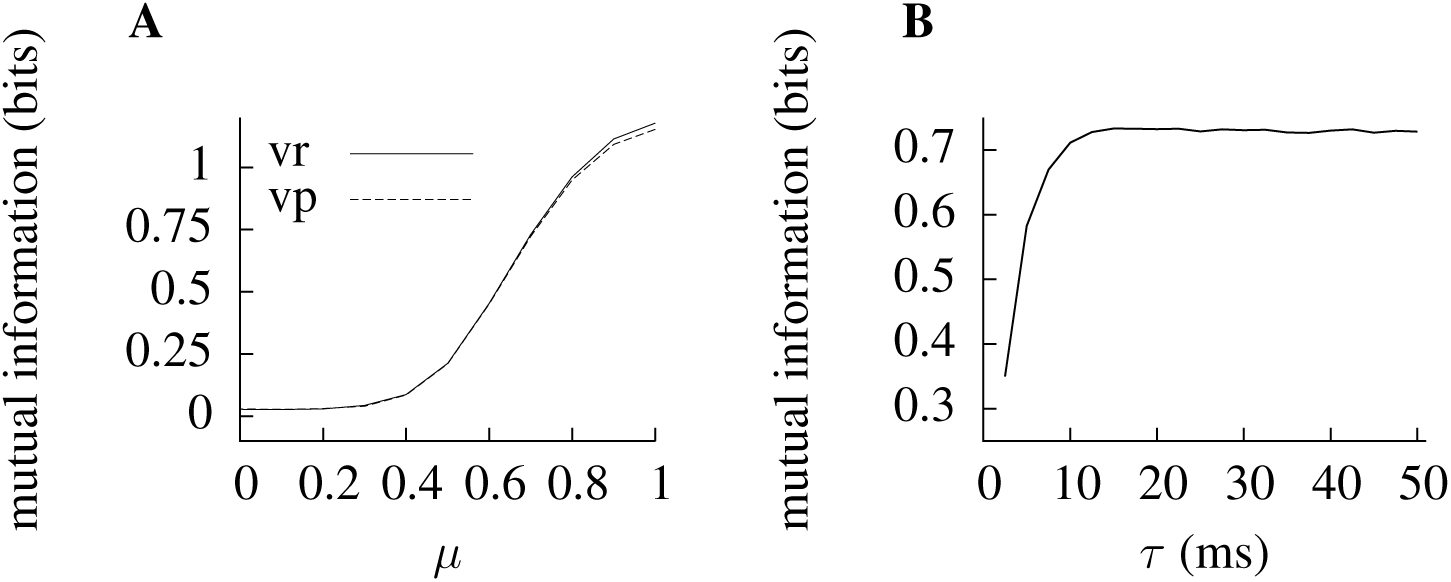
The effect of the metric. In **A** the density estimation approach is applied using the Victor-Purpura metric instead of the van Rossum metric. This comparison uses the same simulated data as Fig. 3, the estimated information using the van Rossum metric with *τ* = 15 ms is marked ‘vr’. The Victor-Purpura metric has a ‘cost’ parameter *q*, like *τ* in the van Rossum metric it expresses the precision of spike times in coding. Here *q* = 2/*τ* and the corresponding estimate is marked ‘vp’. The two estimates are nearly identical. In **B** the value of *τ* used in the metric in the density estimation approach is varied; generally the estimate does not depend sensitively on the value of *τ*.

**Figure 7:**
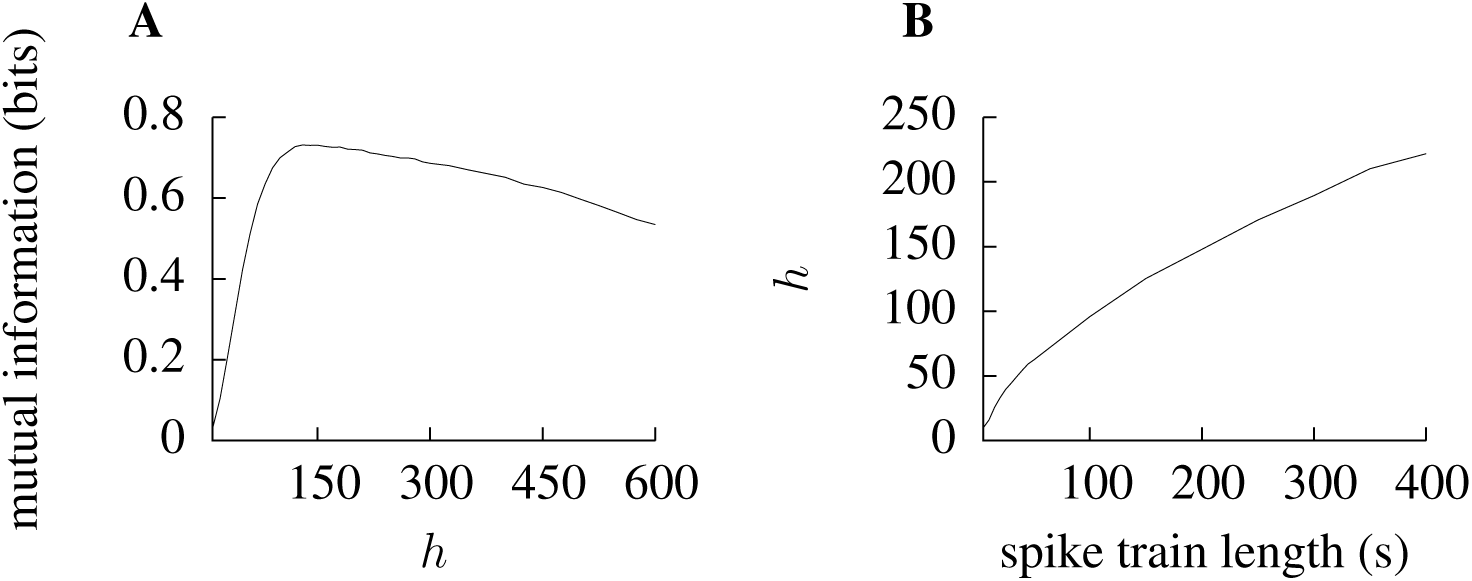
The smoothing parameter *h*. In both these graphs the mutual information is estimated for *μ* = 0.7. In **A** the estimated mutual information is plotted against *h* using 200 s of spike train. In **B** the optimal value of *h*, that is the *h* maximizing the estimated information, is plotted as a function of the spike train length.

## 4 Discussion

This paper describes a method for choosing the smoothing parameter for a Kozachenko-Leonenko estimator of the mutual information and tests it on fictive spike train data. It is seen that the density estimation method is very effective in estimating the mutual information using much smaller amounts of data than that required for the discretization-based approach.

It is hoped that this approach will prove useful in estimating mutual information for real data. Obviously using the approach in practice will require a choice of the interval length and of spike train metric. Typically, as with the binned approach, the interval length should be chosen to reflect the time scale of interest for the cells being studied, this might, for example, reflect the membrane constants of the cells and the correlation length of their stimuli. Hopefully the density estimation method will allow longer interval lengths to be used than was possible with the binned approach. If the van Rossum metric is used, the parameter *τ* needs to be set; ideally this value should maximize the estimated mutual information, often *τ* = 15 ms is used for van Rossum metrics. There are also measures of spike train similarity Kreuz et al. (2007, 2012) that adapt to the time-local spike rate and do not have a parameter like the *τ* in the van Rossum metric, or the *q* in the Victor-Purpura metric; these would avoid the need to fix the metric parameters.

The density estimation approach is more computationally demanding than the binned approach; the whole matrix of distances between pairs of interval pairs must be calculated and for each interval pair the nearest *h* other pairs needs to be found.

In the density estimation method, the information is estimated from the matrix of distance values. This means that any information-carrying features of the spike trains that are not captured by the metric will be lost in the estimate of mutual information. Spike train metrics are often evaluated using transmitted information (Victor and Purpura, 1996; Houghton and Victor, 2010), they are, in this sense, designed to capture the information-carrying features. However, mutual information estimated from a distance matrix must underestimate the true value. In the example considered here, this underestimate appears to be small; there may be some indication that the underestimate increases as the number of spikes increases. This might be a more significant for real data where there may be information-carry motifs in spike trains, there are none in the simulated data.

The Kozachenko-Leonenko estimator is a powerful approach to calculating mutual information based on the proximity structure of the data; it is often more efficient than estimators that do not incorporate this structure. It has also been shown in Houghton (2015) that is does not require that there are useful coördinates for the spaces the random variables take their values on. Obviously this is the case when the data of interest is spike train data, but there are likely to be manifold other applications to other data types, including other applications involving neuroscience data, such as calculating the mutual information between spiking responses and a continuous stimulus space, as previously considered in Panzeri et al. (1999).

## Acknowledgements

Thanks to the James S. McDonnell Foundation for support through a Scholar Award in Cognition (JSMF #220020239; https://www.jsmf.org/). This work was carried out using the computational facilities of the Advanced Computing Research Centre, University of Bristol - http://www.bris.ac.uk/acrc/.

## References

Aronov, D., Reich, D. S., Mechler, F., and Victor, J. D. (2003). Neural coding of spatial phase in v1 of the macaque monkey. Journal of Neurophysiology, 89(6):3304–3327.

Aronov, D. and Victor, J. D. (2004). Non-euclidean properties of spike train metric spaces. Physical Review E, 69(6):061905.

Houghton, C. (2015). Calculating mutual information for spike trains and other data with distances but no coordinates. Royal Society Open Science, 2(5):140391.

Houghton, C. and Kreuz, T. (2012). On the efficient calculation of van rossum distances. Network: Computation in Neural Systems, 23(1-2):48–58.

Houghton, C. and Sen, K. (2008). A new multineuron spike train metric. Neural Computation, 20(6):1495–1511.

Houghton, C. and Victor, J. (2010). Measuring representational distances–the spike-train metrics approach. Visual Population Codes–Toward a Common Multivariate Framework for Cell Recording and Functional Imaging, pages 391–416.

Kiefer, J. (1953). Sequential minimax search for a maximum. Proceedings of the American Mathematical Society, 4(3):502–506.

Kozachenko, L. and Leonenko, N. N. (1987). Sample estimate of the entropy of a random vector. Problemy Peredachi Informatsii, 23(2):9–16.

Kraskov, A., Stögbauer, H., and Grassberger, P. (2004). Estimating mutual information. Physical Review E, 69(6):066138.

Kreuz, T., Chicharro, D., Houghton, C., Andrzejak, R. G., and Mormann, F. (2012). Monitoring spike train synchrony. Journal of Neurophysiology, 109(5):1457–1472.

Kreuz, T., Haas, J. S., Morelli, A., Abarbanel, H. D., and Politi, A. (2007). Measuring spike train synchrony. Journal of neuroscience methods, 165(1):151–161.

Magri, C., Whittingstall, K., Singh, V., Logothetis, N. K., and Panzeri, S. (2009). A toolbox for the fast information analysis of multiple-site LFP, EEG and spike train recordings. BMC Neuroscience, 10(1):81.

Montemurro, M. A., Senatore, R., and Panzeri, S. (2007). Tight data-robust bounds to mutual information combining shuffling and model selection techniques. Neural Computation, 19(11):2913–2957.

Nemenman, I., Bialek, W., and van Steveninck, R. d. R. (2004). Entropy and information in neural spike trains: Progress on the sampling problem. Physical Review E, 69(5):056111.

Nirenberg, S., Carcieri, S. M., Jacobs, A. L., and Latham, P. E. (2001). Retinal ganglion cells act largely as independent encoders. Nature, 411(6838):698.

Panzeri, S., Senatore, R., Montemurro, M. A., and Petersen, R. S. (2007). Correcting for the sampling bias problem in spike train information measures. Journal of Neurophysiology, 98(3):1064–1072.

Panzeri, S., Treves, A., Schultz, S., and Rolls, E. T. (1999). On decoding the responses of a population of neurons from short time windows. Neural Computation, 11(7):1553–1577.

Rossum, M. v. (2001). A novel spike distance. Neural Computation, 13(4):751–763.

Strong, S. P., Koberle, R., van Steveninck, R. R. R., and Bialek, W. (1998). Entropy and information in neural spike trains. Physical Review Letters, 80(1):197.

Tobin, R. J. and Houghton, C. J. (2013). A kernel-based calculation of information on a metric space. Entropy, 15(10):4540–4552.

Treves, A. and Panzeri, S. (1995). The upward bias in measures of information derived from limited data samples. Neural Computation, 7(2):399–407.

Victor, J. D. (2002). Binless strategies for estimation of information from neural data. Physical Review E, 66(5):051903.

Victor, J. D. and Purpura, K. P. (1996). Nature and precision of temporal coding in visual cortex: a metric-space analysis. Journal of Neurophysiology, 76(2):1310–1326.

